# Morphological Alternations of Intraepithelial and Stromal Telocytes in Response to Salinity Challenges

**DOI:** 10.1101/115881

**Authors:** Soha Mohamed Abdel-latief Soliman, Walaa Fathy Ali Emeish

## Abstract

**Summary statement:** The article represent an experimental study in which we investigated the effect of the ssalinty on the communicating cells (telocytes) and their target cells including chloride, stem, Rodlet cells, myoblasts

**Abstract:** Telocyte is a communicating cell established relations to various types of cells. Few experimental studies are performed on telocytes. The current study investigated responce of telocytes to salinity stress in relations to osmoregualtory, immune and stem cells. We exposed Common carp to salinity level 0.2, 6, 10, 14 ppt. Gill samples were fixed and processed for microscopic and TEM. Two types of telocytes were identified: intraepithelial and stromal telocytes. Intraepithelial telocytes comprised the cellular lining of the lymph spaces where they shed the secretory vesicles. Stromal telocytes shed their secretory vesicles in the secondary circulatory vessels. Telocyte enlarged and exhibited high secretory activities. They exert their effect either by direct contact or by paracrine mode. In sanity treated samples, chloride cells enlarged and the mitochondria became cigar-shaped. pavement cells enlarged and micro-ridges elongated. Stromal telocytes established contact with stem cell and skeletal myoblast. Macrophages and Rodlet cells increased in number. In conclusion, intraepithelial and stromal responded to salinity stress by activation of cellular signaling. They play a major role in osmoregulation, immunity, and regeneration.

## introduction

Telocyte is a distinctive type of interstitial cells, which have a wide range of biological functions in different tissues and organs. Functional diversity of telocytes is regarding affecting different types of cells and structures (Varga, Danisovic et al. 2016). Telocytes have unique morphological characteristics. Multiple cell prolongations; telopodes emerge from the cell body and may extend to hundreds of microns. Telopodes may give rise dichotomous branches and establish cellular connections to form a complex labyrinthine system. Telopodes composed of thin segments; podomers and interval expansions; podoms which are rich in calcium release units; mitochondria, endoplasmic reticulum, and caveolae. Telocytes establish a synaptic junction connecting to immunoreactive cells (Popescu and Faussone-Pellegrini 2010).

Telocytes exert their effect on cells either by establishing cellular contact or through paracrine mode. Two types of cellular contact are documented for telocytes; homocellular and heterocellaular contact. Homocellular contact is formed between two telopodes or telocytes and telopodes or between the cell body of two adjacent telocytes. Heterocellular type contact between telocytes and stromal cells either fixed or the free cells. Various types of cellular contacts and communication are mentioned in telocytes including direct apposition of the cell membrane of adjacent telocytes, adherence, and gap junction. Gap junction have a significant role in intercellular signaling pathway (Mirancea 2016). The secretory function of telocytes influence the target cells. Telocytes deliver microvesicles to the cell and providing macromolecules such as proteins or RNAs, microRNA. Telocytes also shed exosomes, ectosomes and multivesicular vesicles (Popescu, Gherghiceanu et al. 2005; Popescu and Faussone-Pellegrini 2010; Cantarero Carmona, Luesma Bartolome et al. 2011)

Telocytes are multifunctional cell functions. They contribute to generation and transmission of nerve impulses to involuntary muscles (Takaki 2003; Hutchings, Williams et al. 2009; Gandahi, Chen et al. 2012; Drumm, Koh et al. 2014). They serve in mechanoreception and may involve in atrial fibrillation (Gherghiceanu, Hinescu et al. 2008). Telocytes exhibit receptors for excitatory and inhibitory neurotransmitters (Iino and Horiguchi 2006). They establish contact with immunoreactive cells such as eosinophil (Cantarero Carmona, Luesma Bartolome et al. 2011), mast cell, and macrophage (Gherghiceanu and Popescu 2012). Telocytes play a role in regeneration of heart, lung, skeletal muscle, skin, meninges and choroid plexus, eye, liver, uterus, urinary system (Bei, Wang et al. 2015).

Several studies are conducted to study telocytes in human and other mammals but few studies are performed in aquatic species. the current study was conducted using the common carp. The carp belong *to Cyprinidae* family which is commonly known in North America as minnow, while in Eurasia is termed as carp. *Cyprinidae family* are freshwater and are uncommon in brackish water; North America, Africa, and Eurasia (Nelson 2006).

Aquatic species regulate ionic exchange to maintain osmotic balance according to environmental salinity. Several organs are involved in osmoregulation including gills, intestine, kidney, skin, operculum (Marshall and Grosell 2005). Marine inhabitants face great challenges to establish ionic balance. Therefore, the ion transporting cells; chloride cells or ionocytes; participate in the elimination of excess ions in seawater fish, while in freshwater fish, chloride cells contribute in ion absorption (Florkin 2014). IN MARINE FISH, Chloride cells are structurally modified to adopt high salinity levels. Fish exposed to high salinity environment acquire a high proportion of mitochondrial-rich chloride cells (Fielder, Allan et al. 2007). Pervious researches studied the salinity in relation to changes of ionocytes cells. Ionocytes serve in osmoregulation via different types of membranous channels; cystic fibrosis transmembrane regulator (CFTR) anion channel, Na,K,2Cl cotransporter (NKCC) and sodium pump (Na,K-ATPase). CFTR is considered as a membrane protein located on the apical surface of many types of epithelial cells. CFTR is a cyclic AMP-dependent chloride channel, a bicarbonate channel and as a modulator of other ion channels (Derichs 2013). The Na-K-Cl cotransporter (NKCC) is a membrane transport proteins that involved in the active transport of sodium, potassium, and chloride ions across the cell membrane (Russell 2000). Na-K-ATPase is an electrogenic transmembrane enzyme located predominantly on the basolateral surface of the chloride cell and actively transport chloride, rather than sodium across the plasma membrane (Suhail 2010). Pavement cells mostly cover the surface the of the filament and lamellar epithelium. They considered as ion transporting cells; their cell membrane is rich in hydrogen ion channels (Laurent, Goss et al. 1994; Perry and Fryer 1997). In the current study, we focused on the communicating cells; telocytes which influence the large population of stromal, muscular, and epithelial cells. The aim of the present study was to investigate morphological alternations of telocytes subjected to salinity stress and their effect on different types of cells with special reference to the osmoregulatory and immune and stem cells in the common carp.

## Materials and methods

### I-Fish source and transportation

The common carp, *Cyprinus carpio.* was obtained from a private fish farm at El- Dakahlea Government and transported in large water tanks. During transportation, the oxygen level was maintained at 5 mg/l and water tank temperature was 23°C ±3 and pH value at 7.2 – 7.5.

### II- Fish acclimation

Apparently, healthy fingerlings fish measured about the length of 7±2 cm and aged 1 month old. The body weight was 10±2 g, and. Fish were collected and transported to the wet laboratory at Faculty of Veterinary Medicine, South Valley University, Qena, Egypt. Fish were maintained under laboratory conditions during adaptation in running water (salinity = 0.2 ppt) for 3 weeks before conducting the experiments and fed twice daily to ad libitum feed on a commercial floating powdered feed containing 45% protein with a feeding rate of 3% of their body weight.

### III- Aquaria

Fish were originally kept in a re-circulating system in porcelain aquaria (260 *65*70cm) according to the protocol of maintaining bioassay fish as was previously described **(Ellsaesser and Clem, 1986)**. Experiments were conducted in fiberglass aquaria with dimensions of 60×30×40 cm. Dissolved oxygen level was maintained above 5 mg/l while water temperature was kept at 23°C ±3 and pH value at 7.2 – 7.5.

### IV- Salinity exposure

36 acclimated, apparently healthy Common carp, *C. carpio* were selected with a body weight range of 9 – 11 g to serve as the experimental groups. Fish were divided into 12 fiberglass aquaria (60×30×40 Cm) to serve as 4 experimental groups, each group contains 9 fish, and there were 3 replicates for each salinity group. Three groups were gradually subjected to three different salinities until concentrations of 6, 10 and 14 ppt with 2 g/L NaCl increase every two days. The fourth group was reared in freshwater; a dechlorinated tape water of 0.2 ppt salinity level and considered as control group. Water was changed every two days with water that had desired salinities, and aquariums were also cleaned at this time. Salinity was checked and adjusted regularly every two days during a water change. When common carp reached the final desired salinity, fish were allowed to acclimate to the new salinities for a minimum of two weeks before sample collections.

### V- Clinical examination of fish

Fish were observed daily during the course of an experiment for any apparent clinical signs, lesions or mortality. Mortality rate was calculated from the number of dead fish between each sampling period.

### VI- Fish sampling

At the end of the period, nine fish were decapitated in each salinity level. Gill filaments and gill arches of both sides were dissected and fixed in glutaraldehyde (10 mL of 2.5% glutaraldehyde and 90 mL 0.1 M Na-phosphate buffered formalin).

### VII- preparation of resin embedding specimens for semi-thin and ultra-thin sectioning

Fixed samples of gill filaments and arches were cut into small pieces. They were washed 4 times for 15 minutes in 0.1 M sodium phosphate buffer (pH 7.2) then were post-fixed in 1% osmic acid in 0.1 M Na-phosphate buffer at 4°C for 2 hours. The osmicated samples were washed 3 times for 20 minutes in 0.1 M phosphate buffer (pH 7.2). Dehydration was performed through graded aceton (70, 80, 90, 100%), 10 minutes for each concentration. The dehydrated samples were immersed in a mixture of aceton/resin (1/1 for 1day, ½ for another day) and pure resin for three days. The resin was prepared by using 10gm ERL, 6gm DER, 26gm NSA and 0.3gm DMAE and thoroughly mixed by a shaker. The specimens were embedded in the resin at 60 C° for 3 days. Polymerized samples were cut to semi-thin sections by using an ultramicrotome Ultracut E (Reichert-Leica, Germany) and stained with toluidine blue (Bancroft, Layton et al. 2013).

Semi-thin sections were also used in histochemical studies. The sections were treated with a saturated alcoholic solution of sodium hydroxide for 15 minutes to dissolve the resin (Lloyd 2001). The semi-thin sections were stained by Heidenhain’s Iron-Hx (Heidenhai 1896) and methylene blue used for staining of paraffin sections and prepared as a stain for semi-thin sections (Bancroft, Layton et al. 2013).

Ultrathin sections were obtained by a Reichert ultra-microtome. The sections (70 nm) were stained with uranyle acetate and lead citrate (Reynolds, 1963) and examined by JEOL100CX II transmission electron microscope (TEM) at the CENTREAL LABARTORY UNIT of South Valley University.

### VIII- Coloring images

Transmission electron microscopy images were colored using photo filter 6.3.2 program. Coloring images required to change the color balance, using the stamp tool to color the objective cells.

## Results

The current study was performed to evaluate ranges of acclimation of the common carp to the hypertonic conditions in relation to responses of the communicating cells; telocytes and the related effector cells including immune, chloride and stem cells in gill filaments and arches using semi-thin and ultrathin sections.

Control and 6ppt salinity exposed groups exhibited normal morphology and behavior and had no noticeable signs of stress and no mortality. However, marked reduction of swimming speed and fish were easily caught in 10 and 14 ppt salinity treated fish.

By semi-thin sections, intraepithelial telocytes had a small cell body and well-defenind telopodes in control samples ( Fig. 1A, E, I). They were gradually enlarged in size during exposure to different levels of salinity. In 6 ppt, intraepithelial telocytes were satellite in shape (Fig. 1B, F, J). In 10 ppt and 14 ppt they were large satellite cells with multiple telopodes Fig. 1C, D, G, H, K, L). stromal telocytes were small and had spindle-shaped cell body form which extended fine telopodes in control samples (Fig. 2A, E, I). In 6 ppt salinity concentration, some stromal telocytes were enlarged and telopodes formed a network (Fig. 2 B, F, J). Telopodes formed an extensive network; secretory vesicles were large and could be easily recognized in samples treated with 10 and 14 ppt salinity levels (Fig 2 C, D, G, H, K, L).

**Figure 1:**
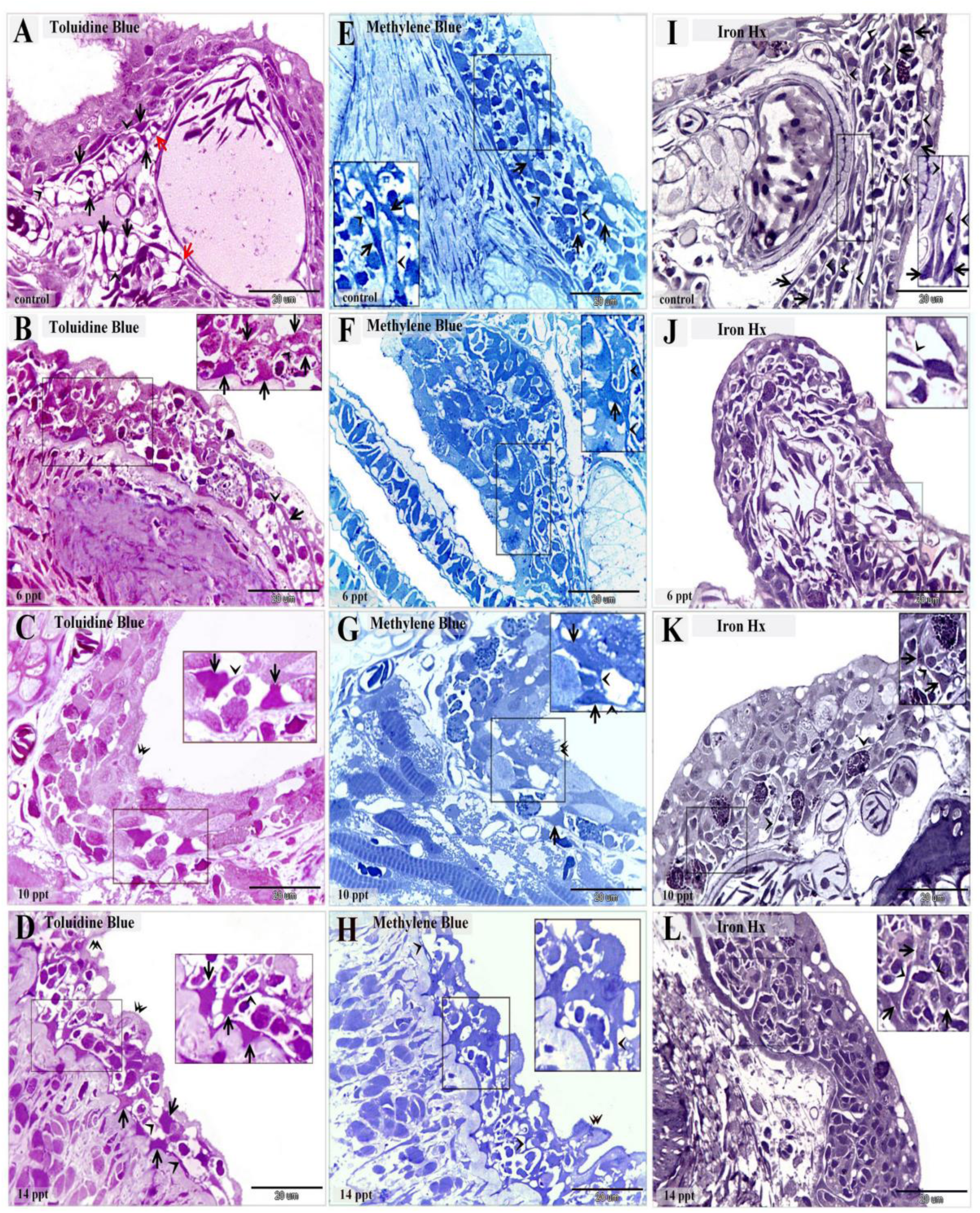
Morphological changes of the intraepithelial telocytes responding to salinity. Semi-thin sections of gill arches and filaments stained with toluidine blue (A-D), methylene blue (E-H) and Heidenhain’s Iron-Hx (I-L). A, E, I: showed intraepithelial telocytes in control samples. They had small cell body (arrows) and prominent telopodes (arrowheads). Note the red arrows refer to the basal lamina. B, F, J: showed intraepithelial telocytes during exposure to 6 ppt salinity level. The cell body enlarged and were satellite in shape (arrows). Note telopodes (arrowheads). C, G, K: cell body undergo hypertrophy and became large satellite (arrows) of the intraepithelial telocytes exposed to 10 ppt salinity level. Note telopodes (arrowheads). D, H, L: the cell body (arrows) of the intraepithelial telocytes increased in size in 14 ppt salinity level. Note telopodes (arrowheads).

**Figure 2:**
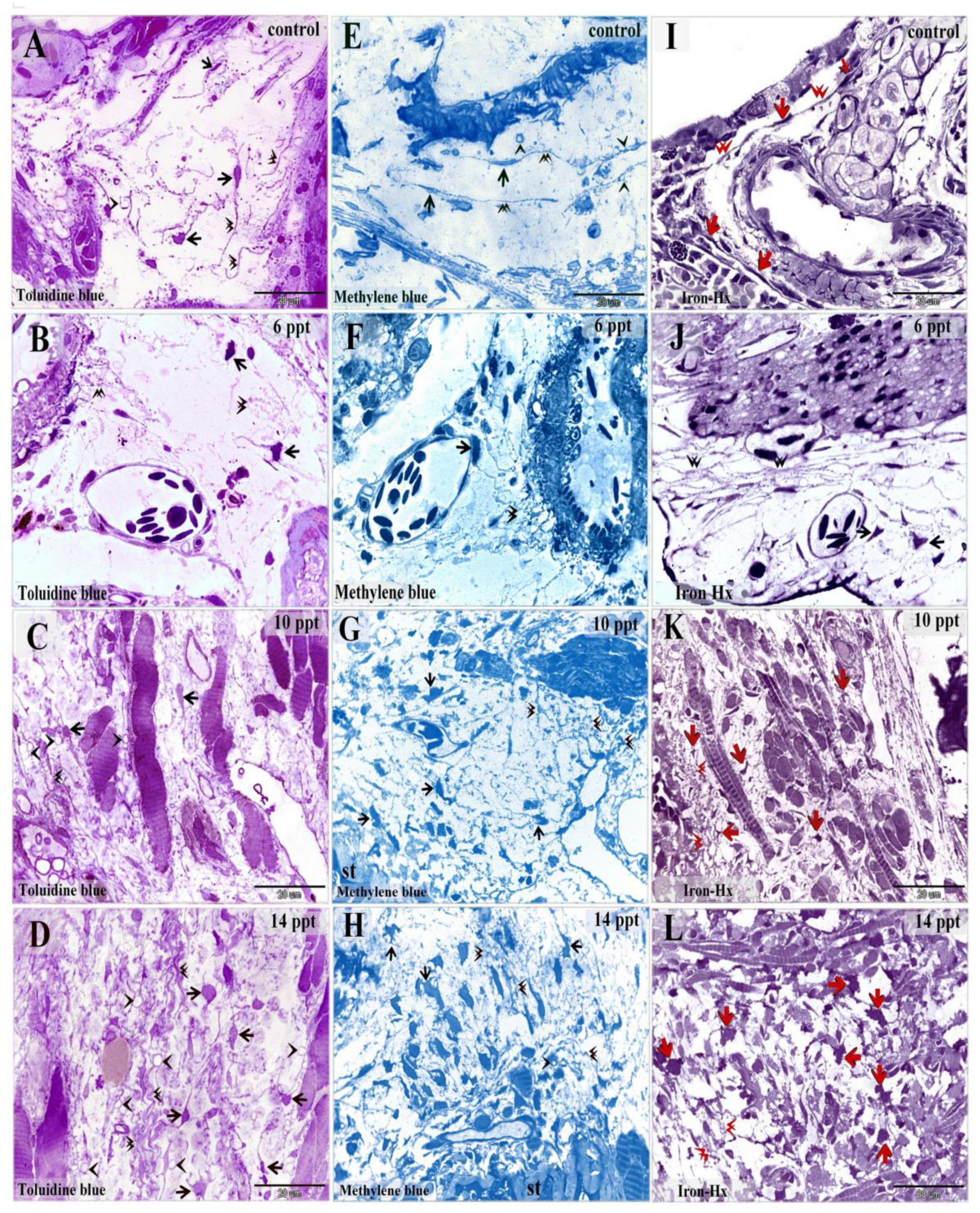
Morphological changes of the stromal telocytes responding to salinity. Semi-thin sections of gill arches and filaments stained with toluidine blue (A-D), methylene blue (E-H) and Heidenhain’s Iron-Hx (I-L). A, E, I: showed stromal telocytes in control samples. They had small cell body (arrows) associated with long telopodes (double arrowheads). Note the secretory vesicles (arrowheads). B, F, J: Some stromal telocytes had enlarged cell body (arrows) during exposure to 6 ppt salinity level. Note telopodes formed a network (double arrowheads). C, G, K: stromal telocytes (arrows) exposed to 10 ppt salinity level had an extensive network of telopodes (double arrowheads) Note the secretory vesicles (arrowheads). D, H, L: stromal telocytes (arrows), telopodes (double arrowheads). Note the secretory vesicles (arrowheads).

**Figure 3:**
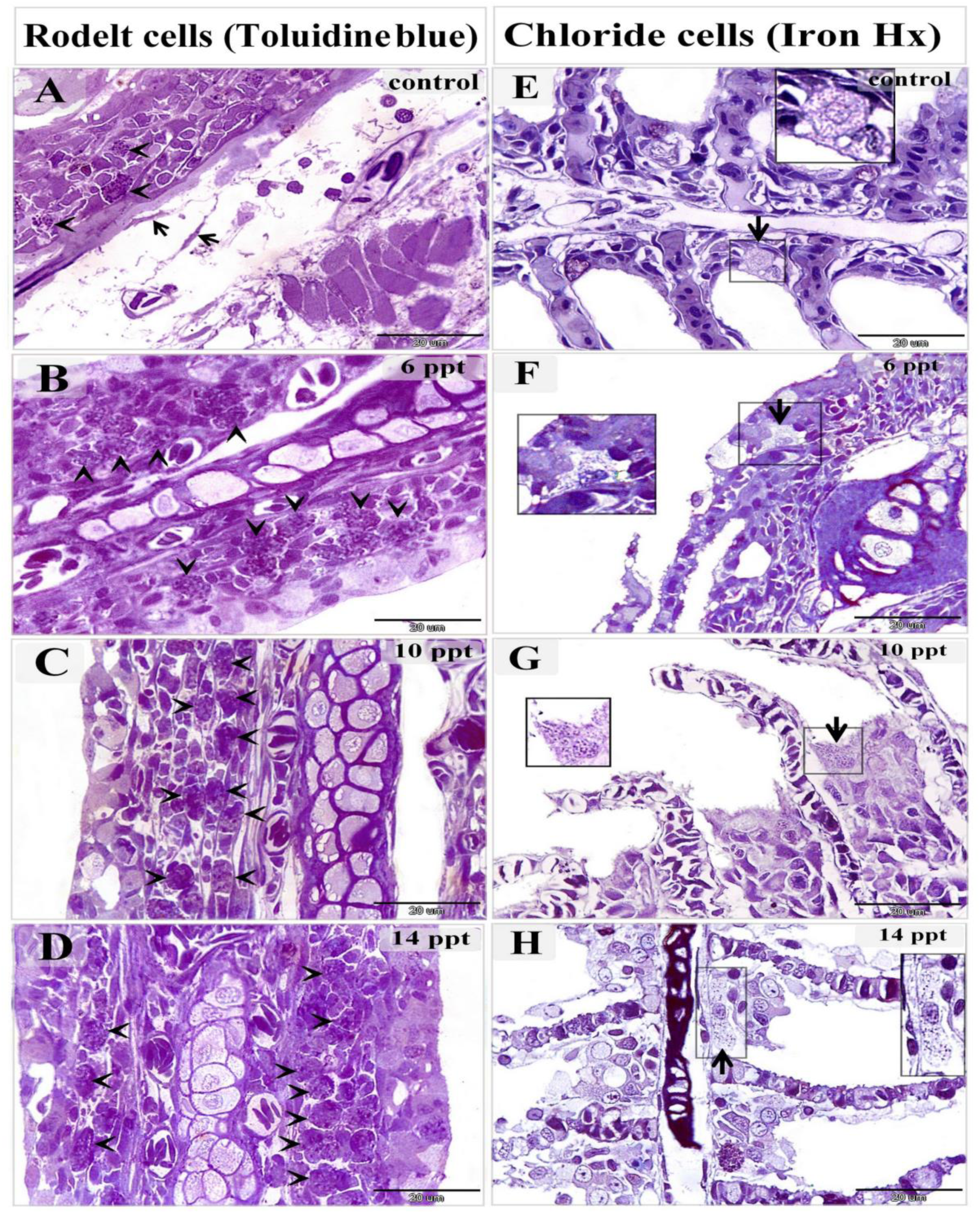
Changes of rodlet and chloride cells responding to salinity stress. Semi-thin sections of gill arches and filaments stained with toluidine blue (A-D), and Heidenhain’s Iron-Hx (E-H). A: few number of immature rodlet cells (granular stage) (arrowheads) in control samples. B: Number of immature rodlet cells (arrowheads) increased in branchial epithelium in 6 ppt treated samples. C: Enormous number of rodlet cells (arrowheads) in branchial epithelium exposed to 10 ppt salinity level. D: Massive number of rodlet cells (arrowheads) in branchial epithelium in 14 ppt treated samples. E: chloride cell ( arrow) in control samples was small contained few mitochondria which appeared as dark dots by iron Hx stain. F: large chloride cells (arrow) in 6 ppt treated samples had larger number of mitochondria. G: Chloride cell (arrow) enlarged and contained abundant mitochondria in 10 ppt treated samples. H: hypertrophy of the chloride cells with massive mitochondrial content in 14 ppt salinity level.

Telocytes were identified for the first time by TEM in the epithelium of the gill arches. In control samples, telocytes represented the cellular lining of the intra-epithelial lymphatic space in which immuno-reactive cells migrate. Intraepithelial telocytes were small, had spindle or satellite-shaped, their telopodes were thin and formed a labyrinth network separating between the compartments of the lymphatic space. Intraepithelial telocytes rest on the basement membrane. Telopodes extended between epithelial and immune cells. Intraepithelial telocytes could establish contact with epithelial cells. The secretory vesicles of the telopodes were excreted in the intraepithelial lymphatic space (Fig 4 A, B). In samples treated with 60 ppt salinity level, intraepithelial telocytes undergo hypertrophy and telopodes were thickened (Fig. 4 C-H).

**Figure 4:**
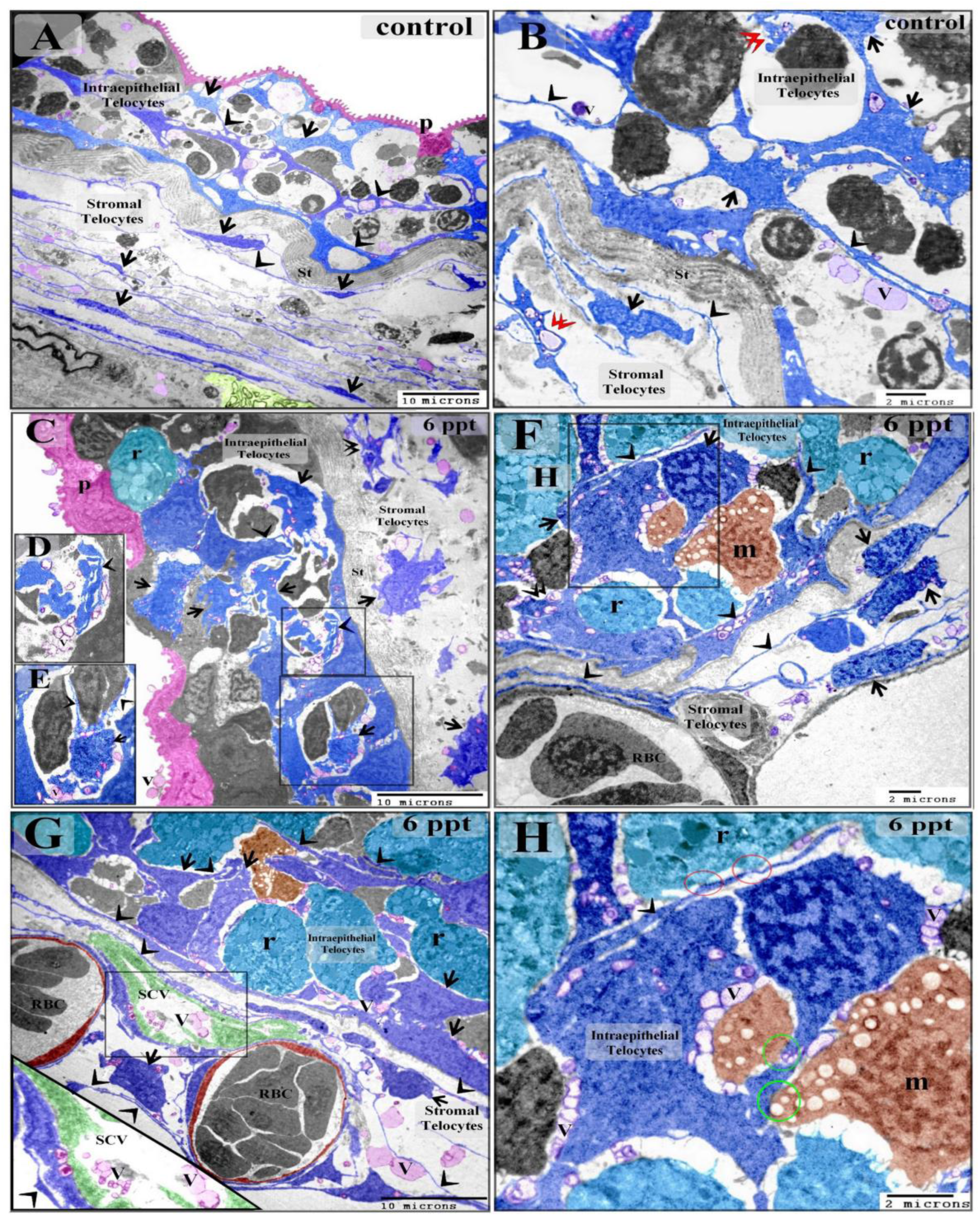
effect of low level (6 ppt) of salinity on intraepithelial telocytes. Colored ultra-thin sections in gill arches (A-F, H) and filaments (G) of control (A, B) and 6 ppt treated samples (C-H). A, B: intraepithelial telocytes were small, had spindle or satellite-shaped (arrows) and thin telopodes (arrowheads). They rest on the basal lamina which directly opposed on the stratum compactum (st). They organized in a labyrinth network which comprised the wall of the epithelial lymph spaces. The secretory vesicles (V) were liberated in the lymph spaces. Note podoms (double arrowheads), pavement cells (P). stromal telocytes had longer and thinner telopodes. C-H: Enlargement of the both intraepithelial and stromal telocytes (arrows), thickening of the telopodes (arrowheads). intraepithelial telocytes liberated the secretory vesicles in the lymph spaces and stromal telocytes shed their vesicles in the secondary circulatory vessels (SCV) or lymphatic vessels. Note stratum compactum (st), pavement cells (p). branchial blood vessels contained red blood cells (RBC). Rodlet cells (r) and lysosome-rich macrophages (m) in the lymph spaces. red circles refer to point of contact between telopodes and rodlet cells, green circle refer to contact between telocytes and the macrophage in branchial epithelium.

In 10 salinity concentration, intraepithelial telocytes was hypertrophy associated with enlargement of the podoms. They also acquired high secretory activity. Intraepithelial telocytes established planar contact with chloride cell (Fig 5A-E). In 14 ppt salinity level, telocytes shed secretory vesicles, exosomes, and multivesicular vesicle into the intra-epithelial lymphatic spaces. Intra-epithelial telocytes established planar contact with chloride cells (Fig 6A- D).

**Figure 5:**
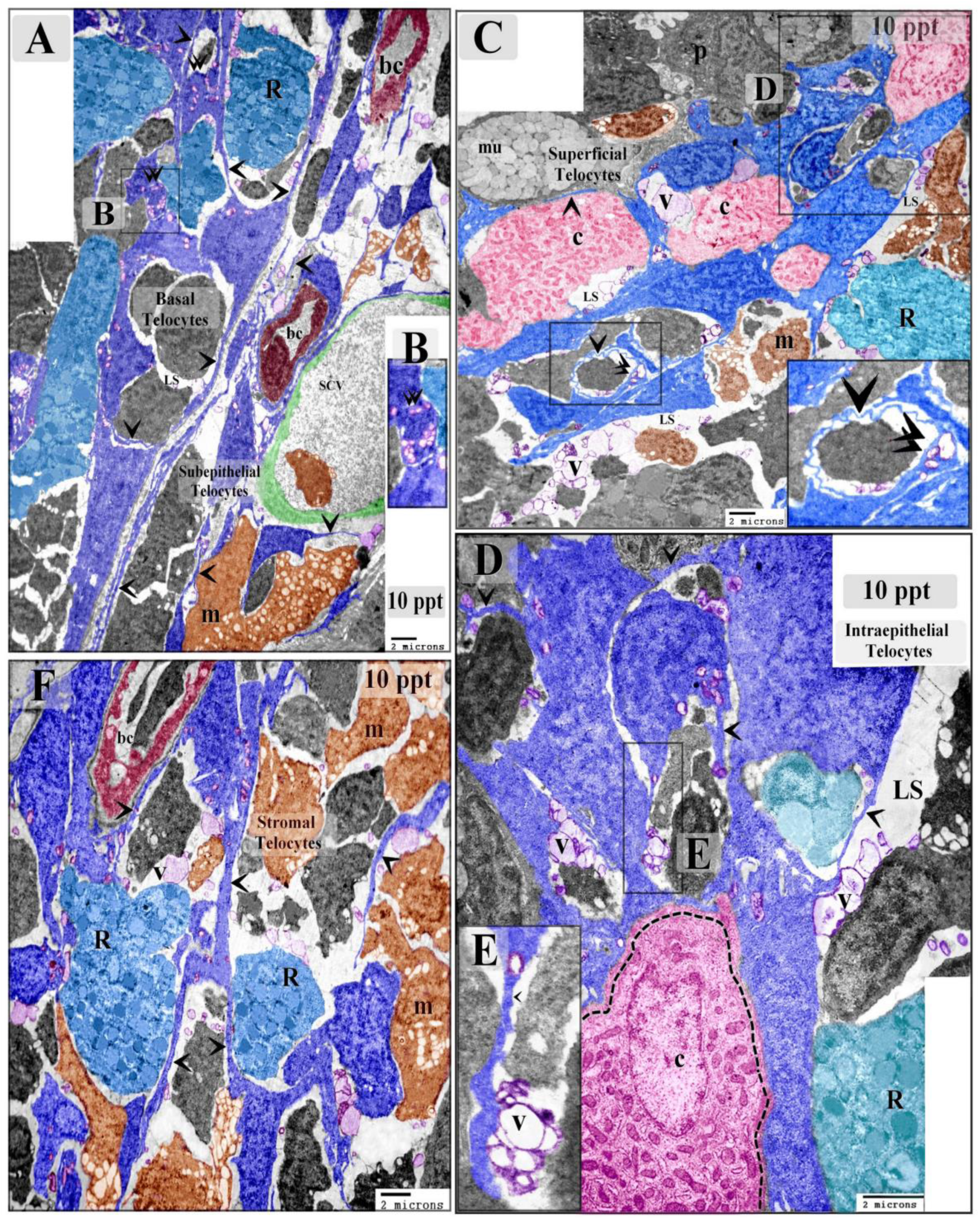
effect of 10 ppt salinity level on intraepithelial telocytes. Colored ultra-thin sections in gill arches of 10 ppt treated samples. The prominent feature in 10 ppt level treated samples was increase in size of telocytes either intraepithelial or stromal. A: Basal telocytes were telocytes in the basal layer of the branchial epithelium established a communicating network between the lymph spaces (LS) in which rodlet cells (R) migrated. Note telopodes (arrowheads), podom (double arrowheads). The subepithelial telocytes connected with blood capillaries (bc), secondary circulatory vessels (SCV) and lysosome-rich macrophages (m). Both Basal intraepithelial and sub-epithelial telocytes undergo hypertrophy. B: high magnification of the podom. C: Superficial intraepithelial telocytes established contact with different types of epithelial cells and formed the lymph spaces (LS) where they shed the secretory vesicles (V). Note telopodes (arrowheads), podoms (double arrowheads). chloride cell (c), rodlet cells (R), pavement cell (P), Mucous cell (mu), macrophages (m). D, E: intraepithelial telocytes established planar contact with chloride cell (dashed line). Telocytes shed the numerous secretory vesicles (V) in the lymph space (LS). Note telopodes (arrowheads). F: Stromal telocytes formed a network connected with different types of stromal cells including rodlet cells (R) and macrophages (m). note telopodes (arrowheads), blood capillary (bc).

**Figure 6:**
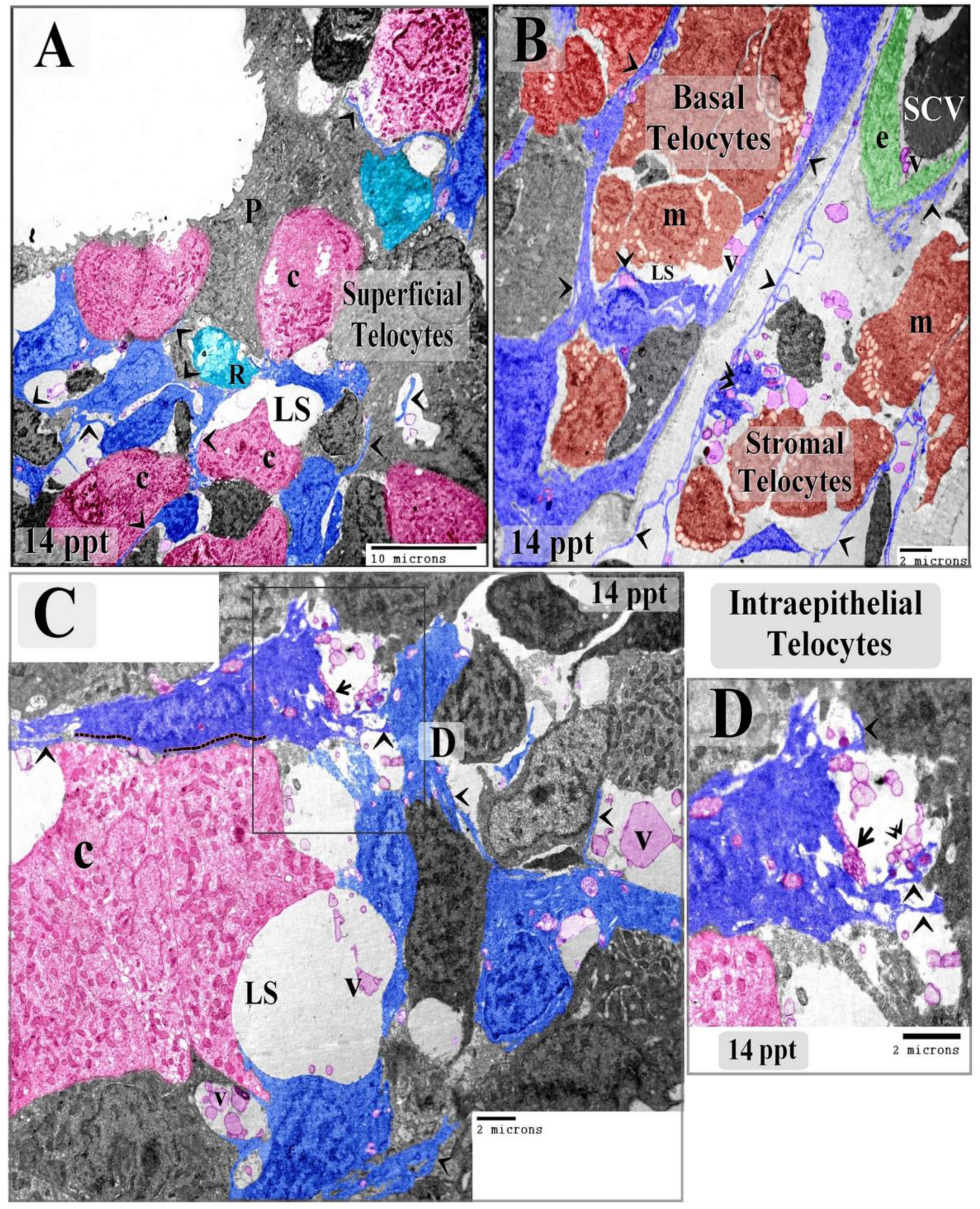
effect of high salinity level (14 ppt) on intraepithelial telocytes. Colored ultra-thin sections in gill arches of 14 ppt treated samples. A: Superficial intraepithelial telocytes communicated forming a labyrinth network between epithelial cells and established the lymph spaces (LS). Note telopodes (arrowheads), Chloride cells (C), rodlet cells (R), pavement cell (P) with short microvilli. B: Basal telocytes organized a network which enclose the lymph spaces (LS). massive number of macrophages (M) in the lymph spaces. Telopodes (arrowheads). Stromal telocytes established contact with secondary circulatory vessels (SCV), stromal macrophages (m). note podom (double arrowhead). C, D: Intraepithelial telocytes formed a network between the epithelial cells and construct the wall of the lymph spaces (LS). Note Intraepithelial telocytes formed a planar contact (dashed line). The secretory vesicles and multivesicular body (arrow), exosomes (double arrowhead) of the intraepithelial telocytes were shed in the lymph spaces. Note telopodes (arrowheads).

Stromal telocytes appeared small spindle, satellite, rounded, triangular shaped cell body with thin cellular prolongations (telopodes) in control samples. Telopodes consisted of podoms and podomers. Macrophages were small and contained vesicles (Fig. 7 A, B, Fig. 8A, B). Telocytes established homocellular junction (Fig 7 C, D). Stromal telocytes undergo morphologically modifications during increasing the concentration of the salinity. The cell body of some populations of telocytes enlarged in salinity levels of 6 ppt (Fig 4 C-H), and 10 ppt (Fig. 5 A, F). In 10 ppt salinity level, telopodes were thickened and became slightly wavy (Fig. 7E, F). Telopodes frequently formed an extensive network (Fig. 8 F). The prominent features of high salinity changes were irregular surface telocyte, waviness, and thickening of telopodes (Fig 7 G, H). They exhibited higher secretory activities in salinity levels reached 6, 10, and 14 ppt (Fig 8 A-D). Stromal telocytes established contact with the endothelial lining of the secondary circulatory system or lymph vessels and liberated the secretory vesicles in proximity to the vessels. Trans-endothelial transportation of the secretory vesicles was observed in the secondary circulatory vessels. The transferring vesicles were shed in the lumen of the secondary circulatory pathway ( Fig. 4G, Fig. 6E, Fig. 8E, F).

**Figure 7:**
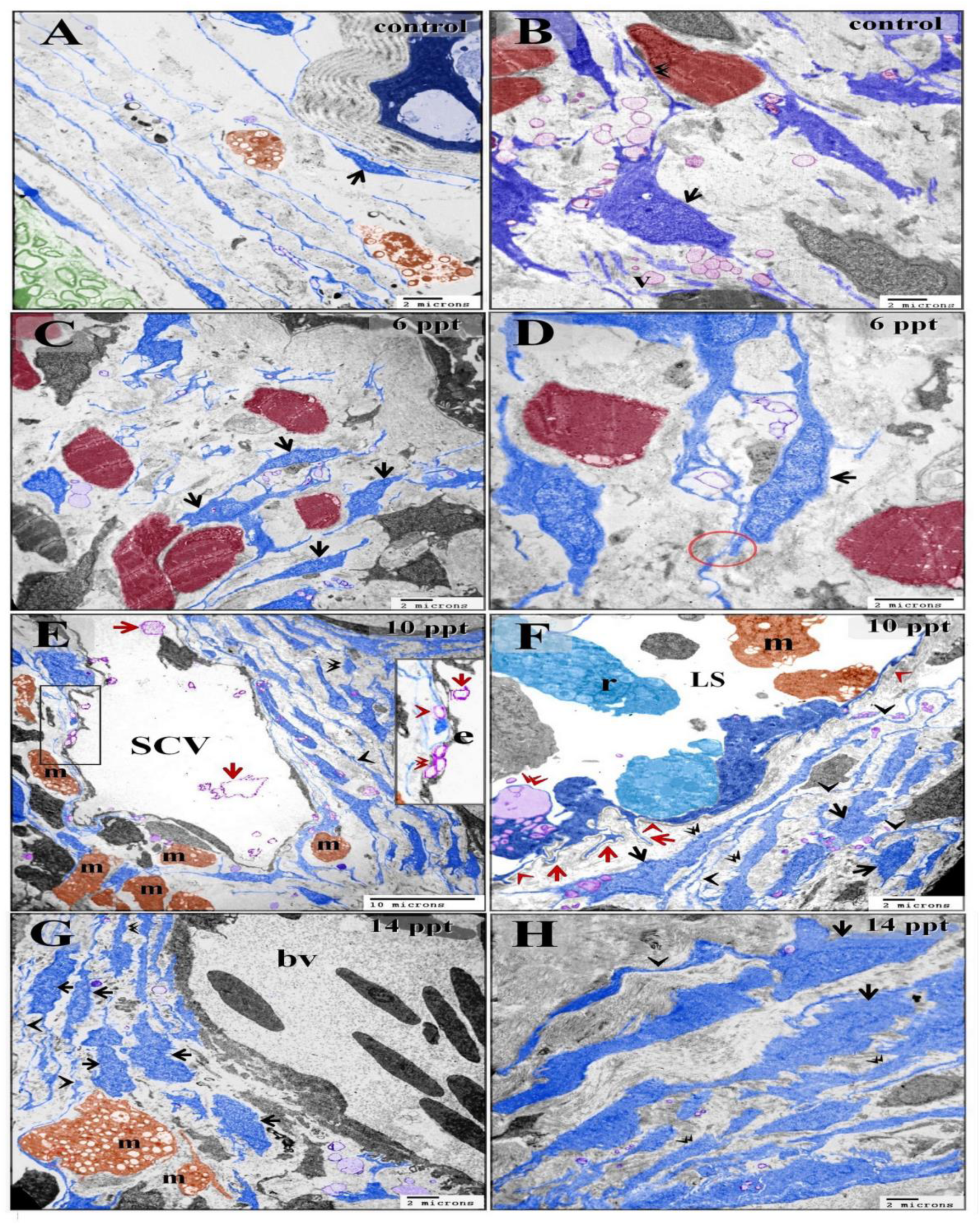
Effect of saintly on stromal telocytes. Colored ultra-thin sections in gill arches of control (A, B), 6 ppt (C, D), 10 ppt (E, F),14 ppt (G, H) treated samples. A, B: telocytes appeared small spindle or satellite shaped (arrows). Telocyte established direct contact with skeletal muscle (double arrowhead). C, D:small spindle-shaped telocytes established homocellular junction (red circle). E: large population of telocytes surrounding the secondary circulatory vessel (SCV). The secretory vesicles (red arrowhead) of telocytes transferred through the endothelial lining of the secondary circulatory vessel (red double arrowheads) and shed in the lumen of the vessel (red arrows). note thick telopodes ( black double arrowhead) and slight waving (black arrowhead) of the telopodes shape, macrophages (m). F: subepithelial telocytes ( black arrows), telopodes became enormous (black arrowheads) telopodes undergo thickening in certain areas (double arrowheads). The basal lamina (red arrowheads). Note the basal intraepithelial telocytes gave rise short basal telopodes (red arrows). Note podom of the basal intraepithelial telocyte (double arrowhead), lymph space (LS), rodlet cells (r), macrophages (m). G, H: The most prominent features of high salinity changes were the cell body had an irregular surface (arrows) and wavy telopodes (arrowheads) and thickening of telopodes (double arrowhead) note macrophage (m) enlargement and was filled with lysosomes (m), blood vessel (bv).

**Figure 8:**
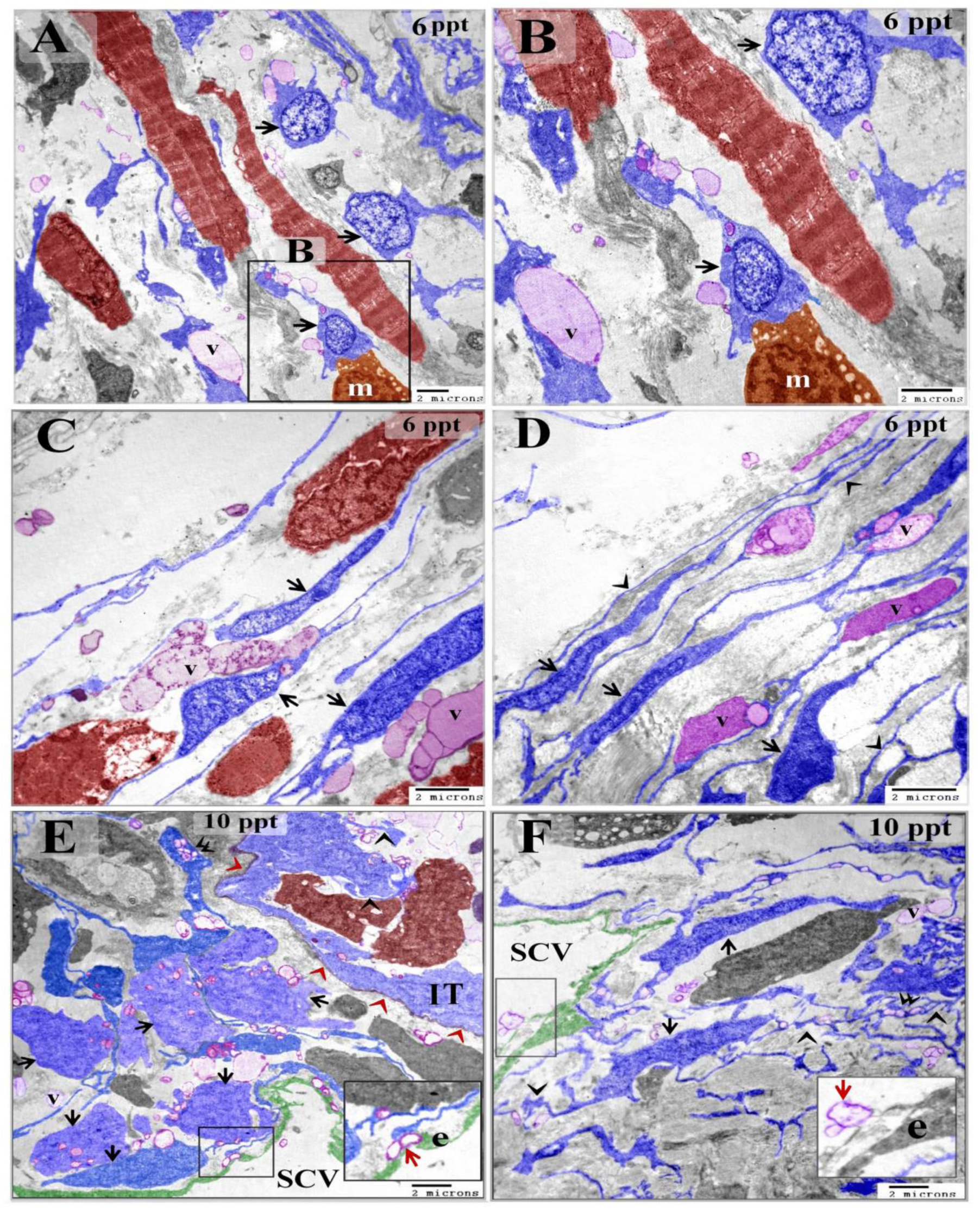
Increase the secretory activities of the stromal telocytes responding to salinity and liberation of the secretory vesicles in the secondary circulatory vessels. Colored ultra-thin sections in gill arches treated with 6 ppt (A-D) and 10 ppt (E, F) level of salinity. A, B: enlarged telocytes acquired rounded or triangular shape (arrows). Note large secretory vesicle (V), cell body of stromal telocyte established contact with macrophage (m). C, D: spindle-shaped telocytes (arrows) shed large secretory vesicle (V). note enormous telopodes (arrowheads). E: large number of enlarged subepithelial telocytes (arrows). Podom (double arrow). Some telocytes established contact with the endothelial lining of the secondary circulatory vessel (SCV). Note transferring of the secretory vesicles (red arrow) secreted by telocytes to the cytoplasm of endothelial cells (e). Basal lamina (red arrowheads). intraepithelial telocytes (IT) and their telopodes (arrowheads). F: spindle shaped stromal telocytes (arrows) and their telopodes formed an extensive network (arrows), podom (double arrowhead), the secretory vesicles (V). note the secretory vesicles (red arrow) were observed in the lumen of the secondary circulatory vessel (SCV). endothelial cells (e).

Telocytes established contact with different types of stromal, epithelial, stem cells and skeletal muscles. Telopodes were connected to nerve fiber (Fig 12A-C) and formed point contact (fig. 7B) or multipoint contact (Fig 12A-C) with skeletal muscles. Large amount of secretory vesicles were excreted closed to the muscular fibers in samples treated with 6 ppt concentration of salinity (Fig 12A, C). An extensive network of telopodes was observed and more secretory vesicles were shed form telocytes. Skeletal muscle fiber undergoes hypertrophy after reaching 10 ppt salinity concentration (Fig. 12D, E).

Both intraepithelial and stromal telocytes established contact with macrophages either via telopodes or the cell body (Fig. 4H, Fig.5 A, F, Fig. 11C, D). In control samples, few macrophages were detected in the gill arches stroma (Fig. 7 A, B). Macrophages became more active and were rich in lysosomes and vesicles in 6 ppt treated samples (Fig. 4 F-H). In 10 ppt salinity level, large number of macrophages in gill arches stroma (Fig. 5 A-F) and epithelial lymphatic spaces (Fig. 5C, Fig. 7E, F). In 14 ppt treated samples, massive lysosomal-rich macrophages were observed in the stroma and epithelial lymphatic spaces (Fig 6B, Fig. 7G).

Telopodes of several telocytes wrapped around stem cells and partially enclosed the stem cell. Telopodes formed a planar contact along the cell membrane of the stem cell, telopodes gave rise small branches extended into the cytoplasm of stem cell (Fig. 11 A, B). Telocytes and their telopodes surrounded and established direct contact with the skeletal myoblast (Fig. 11 C, E, F). Telopodes also formed contact with Schwann cells (Fig. 11 C, D).

Intraepithelial and stromal telocytes formed contact with immature rodlet cells (granular rodlet cells) (Fig. 4F-H, Fig. 5A-D, Fig. 9A, B). Telocytes shed secretory vesicles and multi-vesicular body in the vicinity of immature rodlet cells (Fig. 9A, B). The secretory vesicles of the telocytes were observed in the surface epithelium (Fig. 9 C, D). Intraepithelial telocytes established planar contact with pavement cell (Fig. 9E). In control samples, Pavement cells were flattened with short microvilli (Fig 4A). Pavement cells undergo modifications in salinity treated samples They enlarged in 6 ppt treated samples (Fig. 9C). In 10 ppt salinity concentration, they were cuboidal in shape (Fig. 9E). Pavement cells became elongated and appeared columnar-shaped in 14 ppt level of salinity (Fig. 6A). The micro-ridges became thin, elongated and extended beyond the epithelial surface. Micro-ridges could be seen attached to or enclosing the secretory vesicles of the telocytes. The surface of pavement cells formed pit-like invaginations (Fig. 6 C, D).

**Figure 9:**
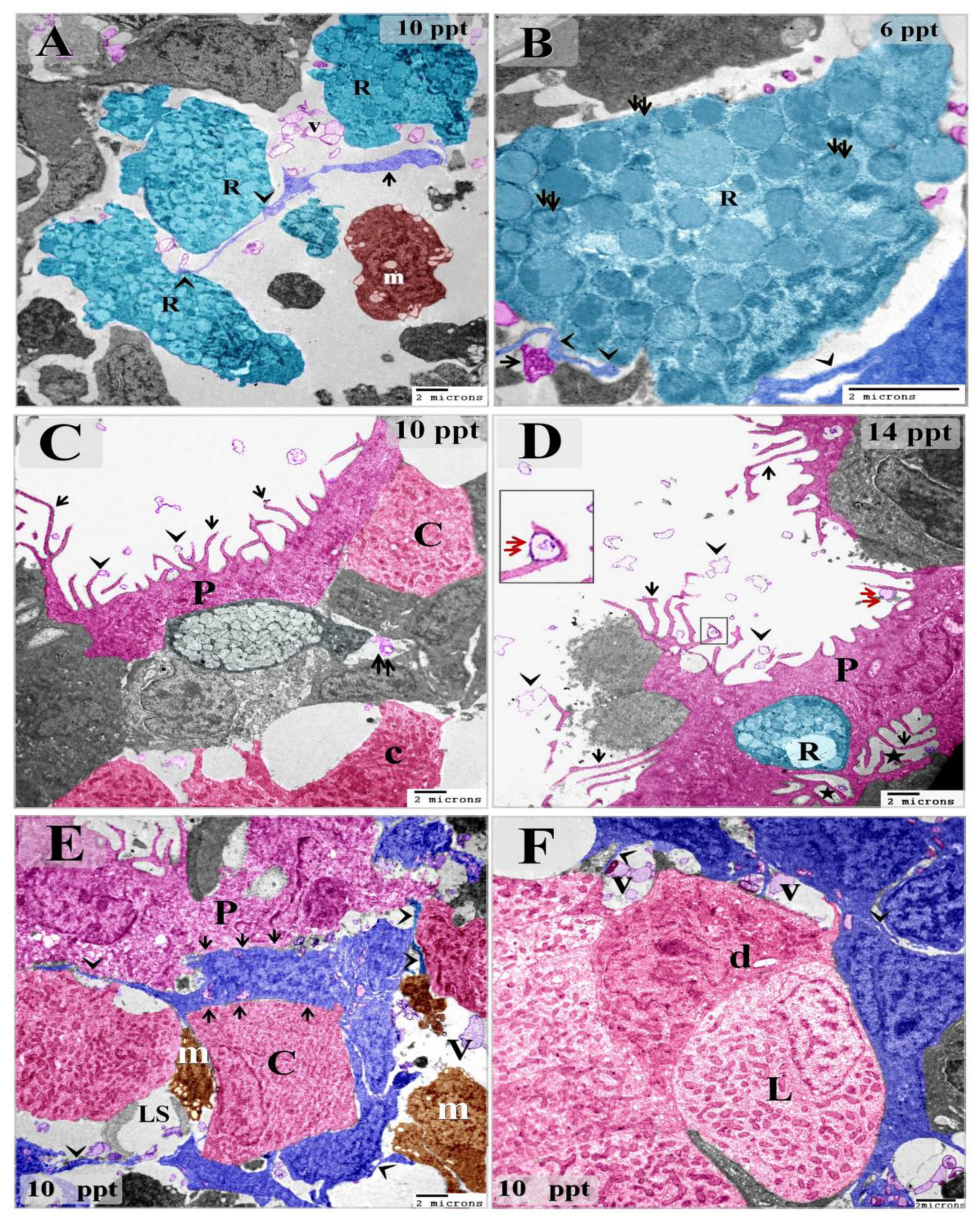
Intraepithelial telocytes in relations to rodlet, chloride and pavement cells. Colored ultra-thin sections in gill arches treated with 6 ppt (B),10 ppt (A, C, E, F) and 14 ppt (D) level of salinity. A: stromal telocytes (arrow) in direct contact with rodlet cells (arrowheads). Note the telocytes shed the secretory vesicles (V) in vicinity to rodlet cells (R). Macrophage (m). B: rodlet cell (granular stage) had immature rodlet granules (double arrows) which contained an electron dense central core. Telopodes (arrowheads) in contact with rodlet cell (R). note multi-vesicular body (arrow). C, D: some pavement cells (P) in 10 and 14 ppt salinity samples had long microvilli (arrows) which deliver the secretory vesicles (arrowheads) of telocytes at the surface of the branchial epithelium. surface invaginations or pits (asterisk) in the pavement cells Note secretory vesicles in lymph space (double arrow), chloride cells (C), rodlet cell (R). E: Intraepithelial telocyte in planer contact (arrows) with the pavement cells (P) and chloride cell (C). note Telopodes (arrows), macrophages (m), secretory vesicles (V), Lymph space (LS). F: two types of mitochondrial rich chloride cells; dark (d) and light (L) were connected to telocytes. Note telopodes (arrows), secretory vesicles (V).

Intraepithelial telocytes established planer contact with Chloride cells (Fig. 5D, 6C). Chloride cells undergo structural modifications during elevation the level of the senility. By TEM, they enlarged, and increase in number gradually depending on salinity concentration. the mitochondrial number increased and changed their morphology from rounded or oval in control group to elongated cigar-shaped in treated samples. Chloride delivered the secretory vesicles of telocytes which were transferred through the intraepithelial lymphatic space (Fig 10 A-D, Fig. 5C, Fig. 6A). Amount of mitochondria in the chloride cells was also evaluated by using Heidenhain’s Iron-Hx. Mitochondria appeared as black granules which increased with the level of the salinity (Fig. 3E-H).

**Figure 10:**
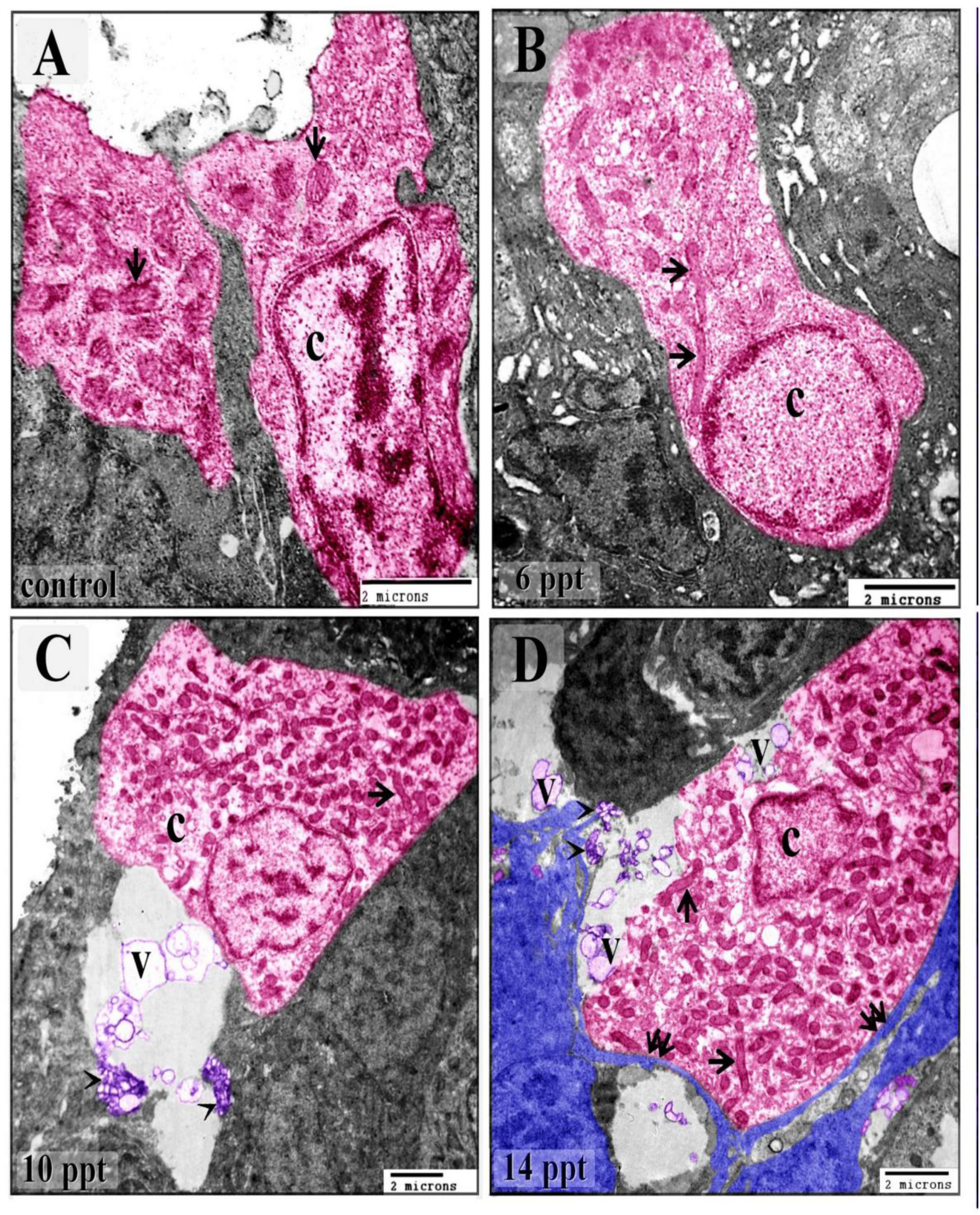
changes of chloride cells in different salinity levels. Colored ultra-thin sections in gill arches control (A) and treated samples with 6 ppt (B),10 ppt (C) and 14 ppt (D) level of salinity. A: chloride cell (C) appeared elongated and had few oval mitochondria (arrows). B: chloride cell (C) increase in size and changed the morphology and appeared more elongated. The mitochondria increased in number and became elongated cigar-shaped (arrows). D: chloride cells (C) enlarged and appeared cuboidal in shape. They had a large number of mitochondria; some of which were elongated cigar-shaped (arrows). Note the secretory vesicles of telocytes in vicinity to chloride cell. E: Chloride cell (C) hypertrophied and appeared oval-shaped. They had a massive mitochondrial content. Some mitochondria were cigar-shaped (arrows). Note telocytes in closed relation to chloride cell. Telopodes (double arrows). The secretory vesicles (V).

**Figure 11:**
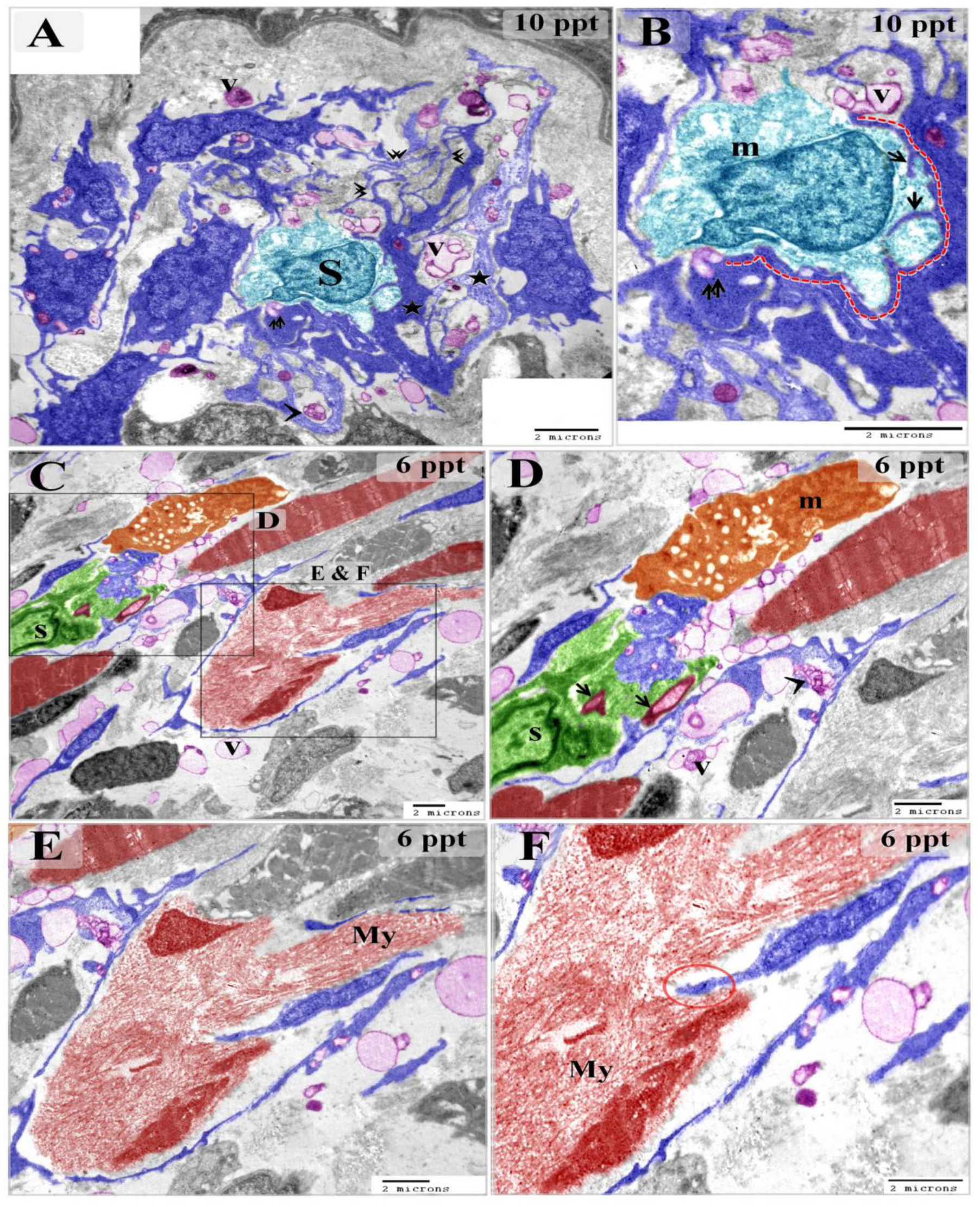
stromal telocytes relation with stem cell and skeletal myoblast. Colored ultra-thin sections in gill arches treated samples with 6 ppt (C-F),10 ppt (A, B) level of salinity. A, B: several stromal telocytes wrapped around stem cell (S) which contained mitochondria (m). telopodes established a planer contact with stem cells (dashed line). telopodes formed an extensive network (double arrowheads). They secreted vesicles (V), multi-vesicular body (arrowhead). Several telopodes interdigitated with the stem cells (arrows). note secretory vesicles attached to stem cell (double arrow). Some telopodes were thickened (asterisk). C-F: telocytes connected with Schwann cell (s) and macrophage (m). note nerve fiber (arrow), secretory vesicles (V), multi-vesicular body (arrowhead). Telocytes established direct contact (red circle) with myoblast which contained ill-organized myofibrils (My).

**Figure 12:**
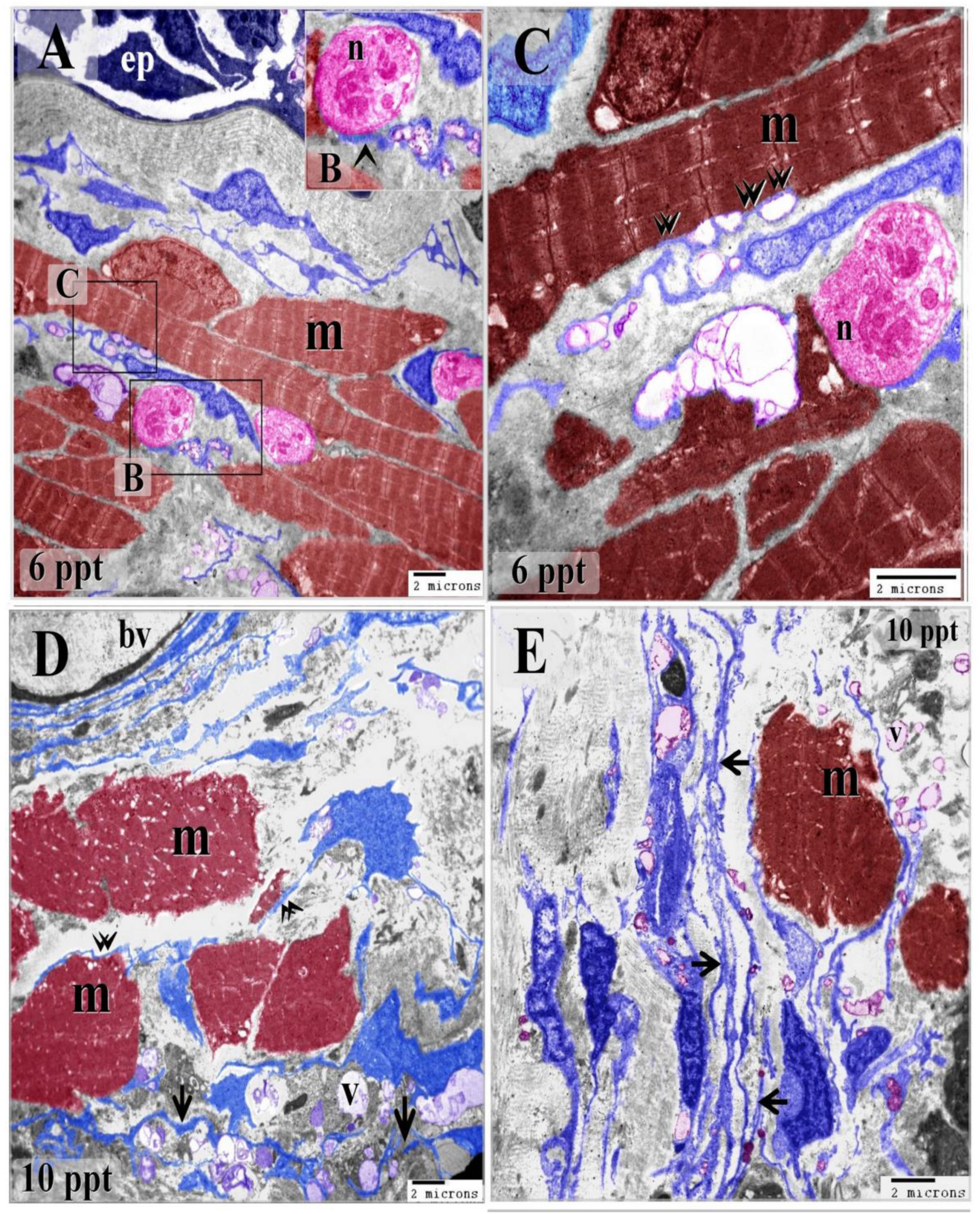
skeletal muscle fibers undergo hypertrophy in response to salinity. Colored ultra-thin sections in gill arches control (A) and treated samples with 6 ppt (A-C),10 ppt (D, E) level of salinity. A, B, C: telocytes established multi-point contact (double arrows) with skeletal muscle fiber (m). telopode formed a direct contact with the nerve fiber (n). note epithelium (ep). D: telocytes established direct contact (double arrowheads) with skeletal muscle fibers increased in diameter (m). note telopodes organized an extensive network (arrows). Note blood vessel (bv), secretory vesicles (V).

## Discussion

The current investigation was carried out to evaluate telocyte response to salinity stress and their relation to osmoregulatory and immune cells. We detected telocytes in semi-thin sections using toluidine blue, methylene blue and Heidenhain’s Iron-Hx, and ultra-thin sections to examine ultrastructural modifications in telocytes in relation to epithelial and stromal cells.

In the current study, telocytes undergo morphological alternations during salinity stress. They were spindle-shaped with fine telopodes in control samples. No significant changes occurred in telocytes in 6 ppt salinity levels, while some telocytes exhibited higher secretory activities. Telocytes shed large secretory vesicles, some populations of telocytes had enlarged cell body, and telopodes became thicker and formed an extensive network, in samples treated with 10 and 14 ppt salinity levels. Hormonal administration could affect the secretory activities of telocytes. Exaggerated secretory activities of telocytes are documented in melatonin treatment of ram seminal vesicles (Abd-Elhafeez, Mokhtar et al. 2016).

In the current study, two types of telocytes were detected in the gills of common carp according to location; intraepithelial telocytes and stromal telocytes. Telopodes formed homocellular and heterocellaular contacts. Heterocellular contact was established with a wide range of cells and structures.

Intraepithelial telocytes communicated to from labyrinth network which comprised the wall of the lymph spaces. These spaces represented interconnected channels interspersed between epithelial cells. Intraepithelial telocytes shed their secretory vesicles and multi-vesicular bodies in the intraepithelial lymph space which deliver them to other epithelial and immune cells including chloride, pavement, mucous, rodlet cells and macrophages. Intraepithelial telocytes may also establish contact with epithelial and immune cells either via point or planar contact. Intraepithelial telocyte is previously detected by scanning electron microscopy in the bovine uterine tube. telocyte is located in the basal layer of the epithelial cells and their telopodes extended between epithelial cells (Abd-Elhafeez and Soliman 2016)

In the current study, intercellular communication between telocytes chloride cells either by direct contact or paracrine mode revealed that telocyte may have a potential role in osmoregulation. Changing salinity level affect telocytes morphology which in turn influence chloride cells. They undergo hypertrophy, change morphology of mitochondria and increase their number upon elevation of the salinity level. similar results are documented in the Hawaiian goby (Stenogobius hawaiiensis). Salinity caused a slight increase in chloride cell number and size (McCormick, Sundell et al. 2003). Gill chloride cells regulate ionic transportation via transport proteins which have a polarized distribution. Three types of transport protein are described in chloride cells; cystic fibrosis transmembrane regulator (CFTR) anion channel, Na,K,2Cl cotransporter (NKCC) and sodium pump (Na,K-ATPase). Expression of Na+/K+-ATPase, Na+/K+/2Cl- cotransporter (NKCC) and cystic fibrosis transmembrane conductance regulator (CFTR) in gill choride cells of the Hawaiian goby (Stenogobius hawaiiensis) is variant in freshwater and 20 per thousand and 30 per thousand salinity concentration for 10 days. Na+/K+-ATPase and NKCC have a basolateral/tubular localization whereas, CFTR expressed in the apical surface of chloride cells. Gill Na+/K+-ATPase expression is not affected by salinity, while CFTR immunoreactivity increase in salinity (McCormick, Sundell et al. 2003).

In the present study, pavement cells were connected with telocytes. The secretory vesicles could reach the surface epithelium and attached or partially enclosed by the micro-ridges of the pavement cells. These cells were flattened in the control samples and gradually enlarged depending on salinity level till became columnar-shaped in 14 ppt salinity concentration. Micro-ridges became thinner and elongated in 10 and 14 salinity levels. Pavement cells developed surface invaginations at 14 ppt salinity. A similar result is described in the gill epithelia of the Adriatic sturgeon Acipenser naccarii. Pavement cells acquired a complex system of microridges on their apical surface during exposure to the hypertonic environment (salinity 35) (CARMONA, GARCIA- GALLEGO et al. 2004). Pavement cells play an important role in gas exchange (Evans, Claiborne et al. 2013). They were rich in proton pumps which regulate acid-based balance (Laurent, Goss et al. 1994; Perry and Fryer 1997).

In the current study, common carp couldn’t sustain salinity levels more than 10 ppt. High mortality rate in fish aquarium began in 12 ppt and was markedly increased in 14 ppt. Mangat and Hundal investigated salinity effect on *Cyprinus carp* survival in different seasons. They used 0, 1.5, 3, 6, and 12 ppt for 60 days. All fish are viable and survive at 0 ppt to 6 ppt salinity during all seasons. Only 50% survive at 12 ppt salinity during winter (14.50C-19.00C), while Fish mortality percentage reached 100% during summer (28.00C-37.00C) and autumn (22.50C-30.50C) (Mangat and Hundal 2014).

In the current study, both intraepithelial and stromal telocytes maintained relation with immune cells; particularly macrophages and rodlet cells; either by cellular contact and paracrine signaling. Thus, telocytes could maintain and enhance immune response in different salinity levels. Macrophages increased in size, number and acquired exaggerated phagocytic activities in samples treated with14 ppt salinity level. They enlarged and became rich in lysosomes and vesicles. Connection of telocytes and macrophage is documented in mouse eye and rat urinary tract (Zheng, Zhu et al. 2012; Luesma, Gherghiceanu et al. 2013). The contact point is identified by an electron-dense nanostructure in the human heart (Gherghiceanu and Popescu 2012). Mouse peritoneal macrophages are activated and secrete macrophage cytokines and enzymes when co-cultured in telocytes conditioned media (Chi, Jiang et al. 2015). Macrophage activity is investigated in relation to salinity level and ration. Phagocytic activities of the macrophages of black sea bream; Mylio macrocephalus Basilewsky juveniles are primarily affected by ration size rather than salinity (Narnaware, Kelly et al. 2001).

In the current study, telocytes formed a planar contact with the stem cell, some telopodes extended into the cytoplasm of stem cell. Moreover, telocytes and their telopodes enclosed and established direct contact with the skeletal progenitor cell which began organization of the intracellular myofilament proteins. Relations between telocytes and stem cells is mentioned in the heart (Popescu, Curici et al. 2015), lung, skeletal muscle, meninges, and choroid plexus (Popescu and Nicolescuy 2013). Cardiac telocytes secrete cytokines and growth factors which promote stem cell proliferation and differentiation such as interleukin (IL)-6, IL-2, IL-10, IL-13, VEGF, macrophage inflammatory protein 1α (MIP-1α), MIP-2 and MCP-1 and some chemokines like, GRO-KC (Albulescu, Tanase et al. 2015).

The present study provided evidence for relations of intraepithelial and stromal telocytes with immature rodlet cells (granular stage). Telocytes exert their effect on rodlet cells either through direct contact or paracrine signaling. Telocytes modified responding to high salinity and subsequently affect rodlet cells. They were increased in number in samples exposed 6, 10 and 14 ppt salinity. Thus, we suggested that telocytes may have a role regulation of the biological activities and in maturation of rodlet cells. Many researches are conducted to investigate the nature and function of rodlet cells. Rodlet cells are thought to act as ion transporting cells and involve in osmoregulation (Ostrander 2000). The widely acceptable hypothesis is that rodlet cells participate in immune response. They are common in helminthic infestations and other noxious agents and considered as a type of eosinophilic granulocyte (Reite and Evensen 2006; Matisz, Goater et al. 2010). Rodlet cells undergo significant changes depending on salinity level. They are increased during reduction of salinity in European sea bass Dicentrarchus labrax (Giari, Manera et al. 2006).

In the present study, stromal telocytes established a direct contact with secondary vascular vessels or lymph vessels. The secretory vesicles of the stromal telocytes were secreted in lymph vessel. Lymphatic vessels deliver the inflow form arterial vessels via arterio-arterial anastomoses and drain into the venous circulation (Kapoor and Bhavna 2004). Thus, we suggested that the secondary vascular vessels represented a principal pathway for trafficking of telocytes sections in the blood circulation. Thus, telocytes may exert their paracrine effect on remote tissues and organs. The secondary vascular vessels are implicated in gaseous exchange and ion transportation (Steffensen and Lomholt 1992).

In conclusion, fish body accommodated changing level of the salinity by activation of an adaptive response through cellular communications. Telocytes represented a major component in the communicating system. They regulated the function of a wide variety of cells either by direct contact or paracrine mode. Morphological modification of telocytes in different levels of salinity reflected increase their activities which influence epithelial, immune and stromal effector cells. Intraepithelial telocytes affected chloride, pavement cells, immature rodlet cells and macrophages while stromal telocytes influence stem cells, skeletal myoblasts and also macrophages and rodlet cells. Thus, telocytes enhanced immunity, osmoregulation in gill lamellar and filament epithelium and regeneration of stromal cells. Thus, the fish could sustain hyperosmotic environments reaching 10 ppt and maintain internal homeostasis.

